# Power and pitfalls of computational methods for inferring clone phylogenies and mutation orders from bulk sequencing data

**DOI:** 10.1101/697318

**Authors:** Sayaka Miura, Tracy Vu, Jiamin Deng, Tiffany Buturla, Jiyeong Choi, Sudhir Kumar

## Abstract

**Background:** Tumors harbor extensive genetic heterogeneity in the form of distinct clone genotypes that arise over time and across different tissues and regions of a cancer patient. Many computational methods produce clone phylogenies from population bulk sequencing data collected from multiple tumor samples. These clone phylogenies are used to infer mutation order and clone origin times during tumor progression, rendering the selection of the appropriate clonal deconvolution method quite critical. Surprisingly, absolute and relative accuracies of these methods in correctly inferring clone phylogenies have not been consistently assessed.

**Methods:** We evaluated the performance of seven computational methods in producing clone phylogenies for simulated datasets in which clones were sampled from multiple sectors of a primary tumor (multi-region) or primary and metastatic tumors in a patient (multi-site). We assessed the accuracy of tested methods metrics in determining the order of mutations and the branching pattern within the reconstructed clone phylogenies.

**Results:** The accuracy of the reconstructed mutation order varied extensively among methods (9% – 44% error). Methods also varied significantly in reconstructing the topologies of clone phylogenies, as 24% – 58% of the inferred clone groupings were incorrect. All the tested methods showed limited ability to identify ancestral clone sequences present in tumor samples correctly. The occurrence of multiple seeding events among tumor sites during metastatic tumor evolution hindered deconvolution of clones for all tested methods.

**Conclusions:** Overall, CloneFinder, MACHINA, and LICHeE showed the highest overall accuracy, but none of the methods performed well for all simulated datasets and conditions.

## Background

Somatic mutations play a crucial role in cancer progression [1–3]. Early models proposed that clones with driver mutations sweep through the population, which is called a linear progression of clone evolution [4]. Now, it is clear that tumors are not monoclonal, and that the clonal evolution generally follows a branching model (i.e., incomplete clonal sweep) even within a tumor [4–10]. Similarly, metastatic tumors also follow a branching pattern [11, 12]. Clones found in primary and metastatic tumors show inter- and intra-tumor evolutionary relationships, which can be represented by a single-patient clone phylogeny [13–16] (e.g., **Fig. 1g and 1h**). The reconstruction and analysis of clone phylogenies have become standard practices in cancer genomics [16–26].

**Figure 1.**
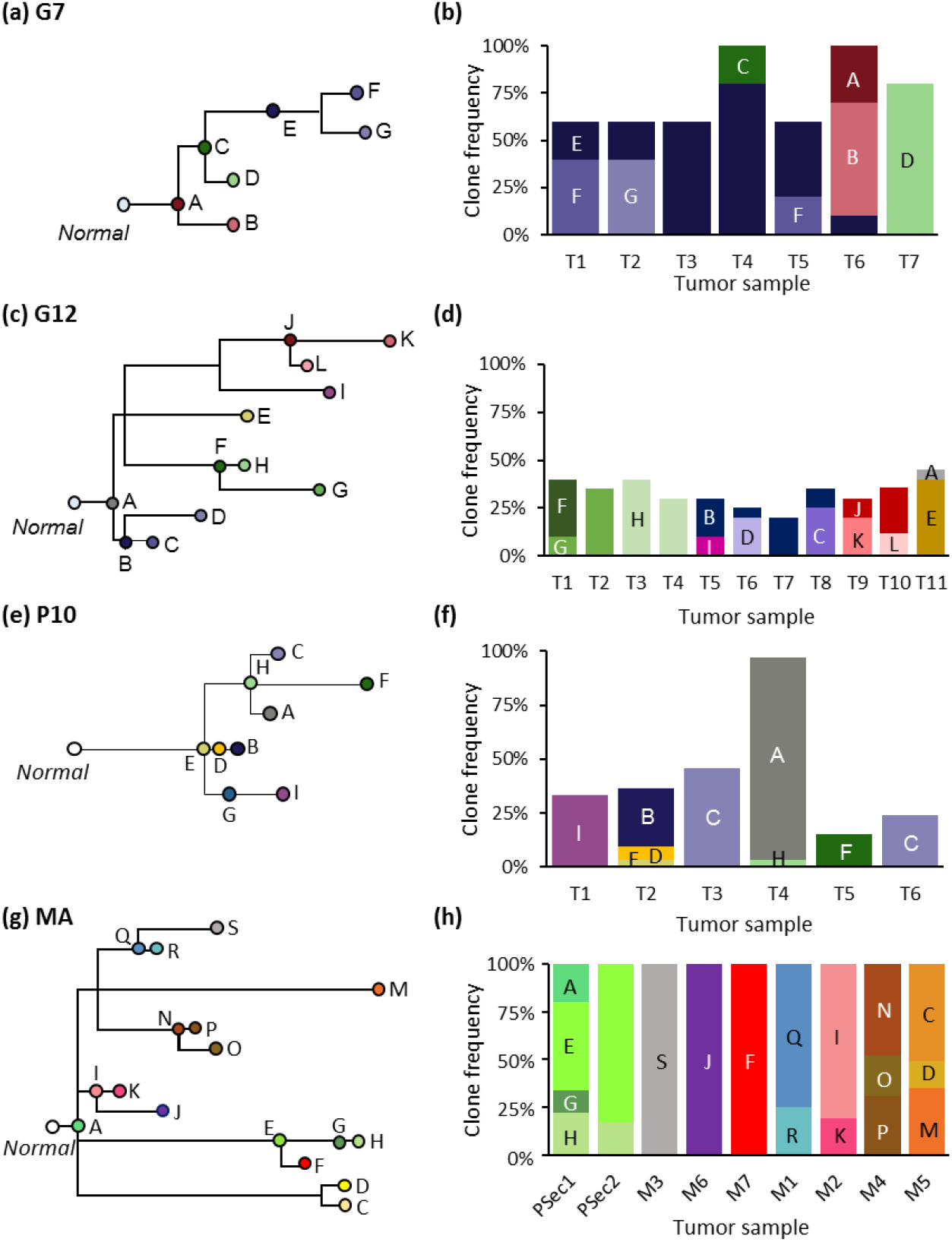
Simulated clone phylogenies and tumor composition. (**a and b**) A phylogeny, and clone frequencies of seven clones and seven tumor samples (T1-T7) derived from EV005 tree (G7 datasets) [44]. (**c** and **d**) A phylogeny and clone frequencies of twelve clones and eleven tumor samples (T1-T11) derived from RK26 tree (G12 datasets) [44]. (**e** and **f**) One of thirty phylogenies and its tumor composition from P10 datasets [35]. (**g** and **h**) One example of MA datasets (out of the 60) with primary tumor (PSec1 and PSec2) and metastatic tumors (M1-M5) [13]. Note that tumor purities are 100% for all the samples.

Clone phylogenies are most often inferred using bulk sequencing data [16, 27–30]. Bulk sequencing of tumor samples is cost effective and can accurately identify single nucleotide variants (SNVs) [31, 32]. The resulting data contains SNV frequencies of cancer cell populations within each tumor sample [27, 33]. Several computational methods have been developed to decompose these SNV profiles into individual clone genotypes, and to predict clone phylogenies [13, 34–39]. These clone genotypes and phylogenies are then employed to infer relative ordering of somatic mutations and to build migration maps of metastatic tumors [40, 41].

Computational methods for clone prediction and phylogeny inference are operationally different from each other. PhyloWGS clusters together SNVs at similar frequencies and then orders them to infer clone genotypes and phylogeny [37]. MACHINA follows a similar process, but also incorporates a model of cancer cell migration between tumor sites (seeding events) [13]. LICHeE generates SNV clusters defined by the pattern of presence and absence of SNVs among tumor samples while considering SNV frequencies [34]. CloneFinder reconstructs ancestral clones in predicting clone genotypes [35]. Treeomics examines the presence and absence of SNVs among tumor samples and resolves evolutionarily incompatible patterns when decomposing SNV profiles into clone genotypes [36]. Ultimately, all of these methods deconvolute individual clones from population bulk sequencing of multiple tumor samples acquired over time and/or different locations in a patient.

Surprisingly, absolute and relative accuracies of clone phylogenies produced by these computational methods have not been assessed using the same collection of datasets, i.e., their performances are yet to be benchmarked. Such benchmarking is critical, because of the biological relevance of the downstream inferences. For example, the accuracies of the order of driver mutations and the interrelationship of clones depend on the performance of current methods in accurately deconvoluting individual clone genotypes and reconstructing evolutionary events. No previous study has evaluated the relative accuracy of clone phylogenetic inferences, as they focused on introducing and assessing the strengths of the new clone prediction methods [13, 34–39]. Besides, the robustness of these computational methods to the complexity of clonal structures and the evolutionary histories of clones from different tumor sites is largely unknown.

Therefore, we evaluated the accuracy of clone phylogenetic inferences by seven clone prediction methods (**Table 1**). We used bulk sequencing datasets simulated under various tumor evolutionary scenarios. Simulated data included small and large numbers of persistent ancestral clones and metastatic tumors that arise from polyclonal seeding events. Our assessments are based on simulation studies because correct phylogenies are known, and computer simulation has emerged as a standard approach for evaluating the performance of statistical methods in cancer genomics [34, 35, 37, 42]. In this study, we identify and highlight the limitations of methods that can most accurately infer clone phylogenies.

**Table 1.**
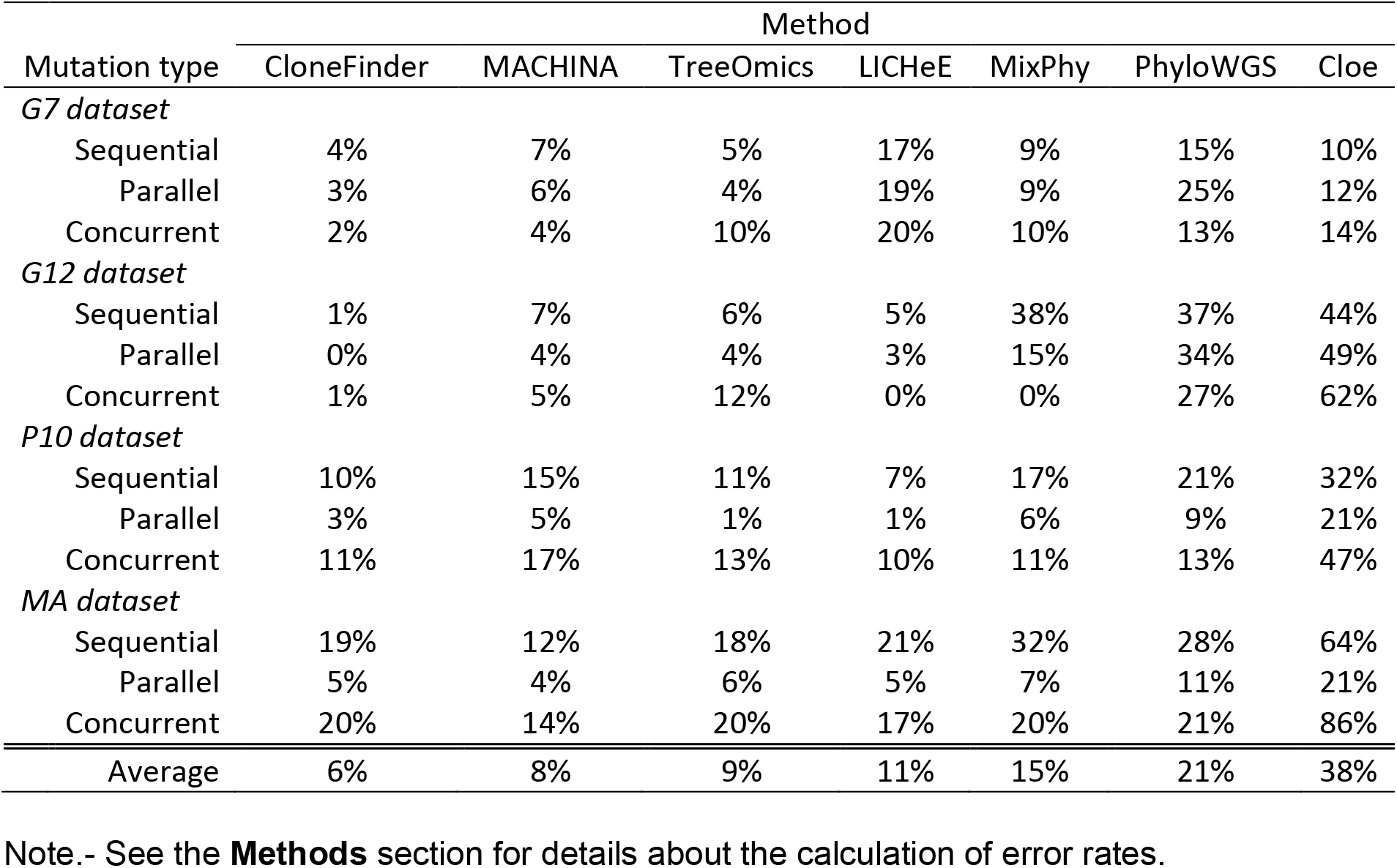
Error rates of methods in inferring sequential, parallel, and concurrent mutations.

| Mutation type | Method |  |  |  |  |  |  |
| --- | --- | --- | --- | --- | --- | --- | --- |
|  | CloneFinder | MACHINA | TreeOmics | LICHeE | MixPhy | PhyloWGS | Cloe |
| *G7 dataset* |  |  |  |  |  |  |  |
| Sequential | 4% | 7% | 5% | 17% | 9% | 15% | 10% |
| Parallel | 3% | 6% | 4% | 19% | 9% | 25% | 12% |
| Concurrent | 2% | 4% | 10% | 20% | 10% | 13% | 14% |
| *G12 dataset* |  |  |  |  |  |  |  |
| Sequential | 1% | 7% | 6% | 5% | 38% | 37% | 44% |
| Parallel | 0% | 4% | 4% | 3% | 15% | 34% | 49% |
| Concurrent | 1% | 5% | 12% | 0% | 0% | 27% | 62% |
| *P10 dataset* |  |  |  |  |  |  |  |
| Sequential | 10% | 15% | 11% | 7% | 17% | 21% | 32% |
| Parallel | 3% | 5% | 1% | 1% | 6% | 9% | 21% |
| Concurrent | 11% | 17% | 13% | 10% | 11% | 13% | 47% |
| *MA dataset* |  |  |  |  |  |  |  |
| Sequential | 19% | 12% | 18% | 21% | 32% | 28% | 64% |
| Parallel | 5% | 4% | 6% | 5% | 7% | 11% | 21% |
| Concurrent | 20% | 14% | 20% | 17% | 20% | 21% | 86% |
| Average | 6% | 8% | 9% | 11% | 15% | 21% | 38% |

Note.- See the **Methods** section for details about the calculation of error rates.

## Results

We analyzed 150 simulated datasets of tumor bulk sequencing data in which the number of tumor samples ranged from 6 to 11. Tumors and clone sequences were simulated with four distinct models of branching evolution (G7, G12, P10, and MA datasets; **Fig. 1**), and a variety of simulated clone phylogenies (e.g., **Fig. 2**). Details of these simulated datasets are described in the **Methods** section. We inferred clone phylogenies for each simulated dataset by using seven different methods (**Table 1**). We used multiple metrics to assess the accuracy, including those measures that score the correctness of the order of mutations and the branching order within the reconstructed clone phylogenies.

**Figure 2.**
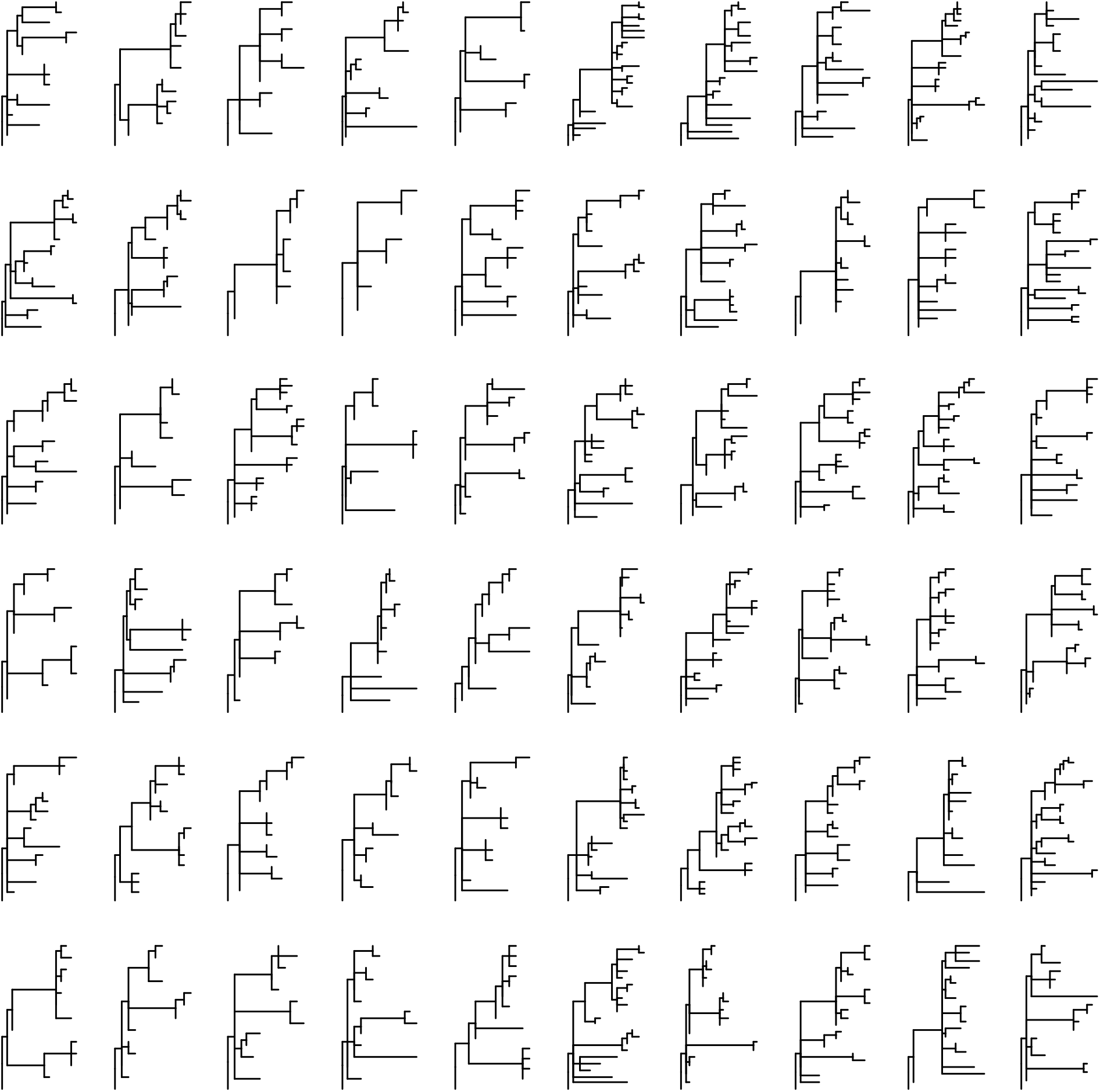
Clone phylogenies used for simulating MA datasets. All clone phylogenies were different.

### Accuracy of ordering mutations

A clone phylogeny can be viewed as a mutational tree [43] in which all the mutations are mapped along branches (e.g., **Fig. 3**). Such mutational trees can be used to test whether a pair of mutations have occurred concurrently, sequentially, or in parallel (**Fig. 3**). At first, we evaluated the accuracy of the predicted order of mutations by using the MLTED score; a smaller score shows greater similarity between the true and inferred mutational tree (see the **Methods** section for details). We begin with results for G7 and G12 datasets that were modeled after the predicted evolutionary histories of two patients (EV005 and RK26, respectively) (**Fig. 1a-1d**) [35, 44]. Each tumor sample may contain one or a few evolutionarily closely-related clones, assuming a localized genetic heterogeneity [4, 6], i.e., migration of cancer cells to another section of a tumor was assumed to be rare. In total, we obtained 60 simulated datasets (replicates) with 34-89 SNVs per dataset. G7 datasets contained seven tumor samples per dataset, while G12 datasets contained eleven samples. For the G7 datasets, all seven methods showed relatively small MLTED scores. For the G12 datasets, four methods (CloneFinder, MACHINA, Treeomics, and LICHeE) produced much smaller MLTED scores compared to other three (PhyloWGS, MixPhy, and Cloe) (**Fig. 4a**).

**Figure 3.**
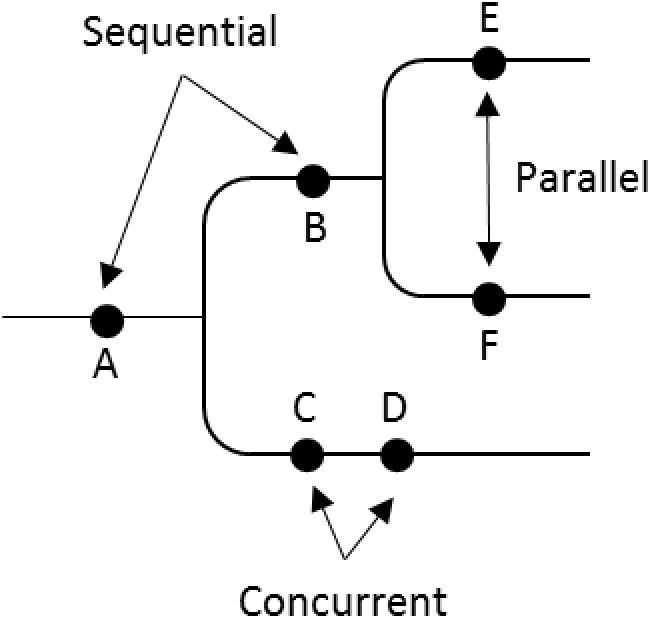
A mutational tree with concurrent (e.g., C and D), sequential (e.g., A and B), and parallel (e.g., E and F) mutations. Dots depict mutations. Order of mutations on a branch (e.g., C and D) cannot be determined based on the clone phylogeny alone.

**Figure 4.**
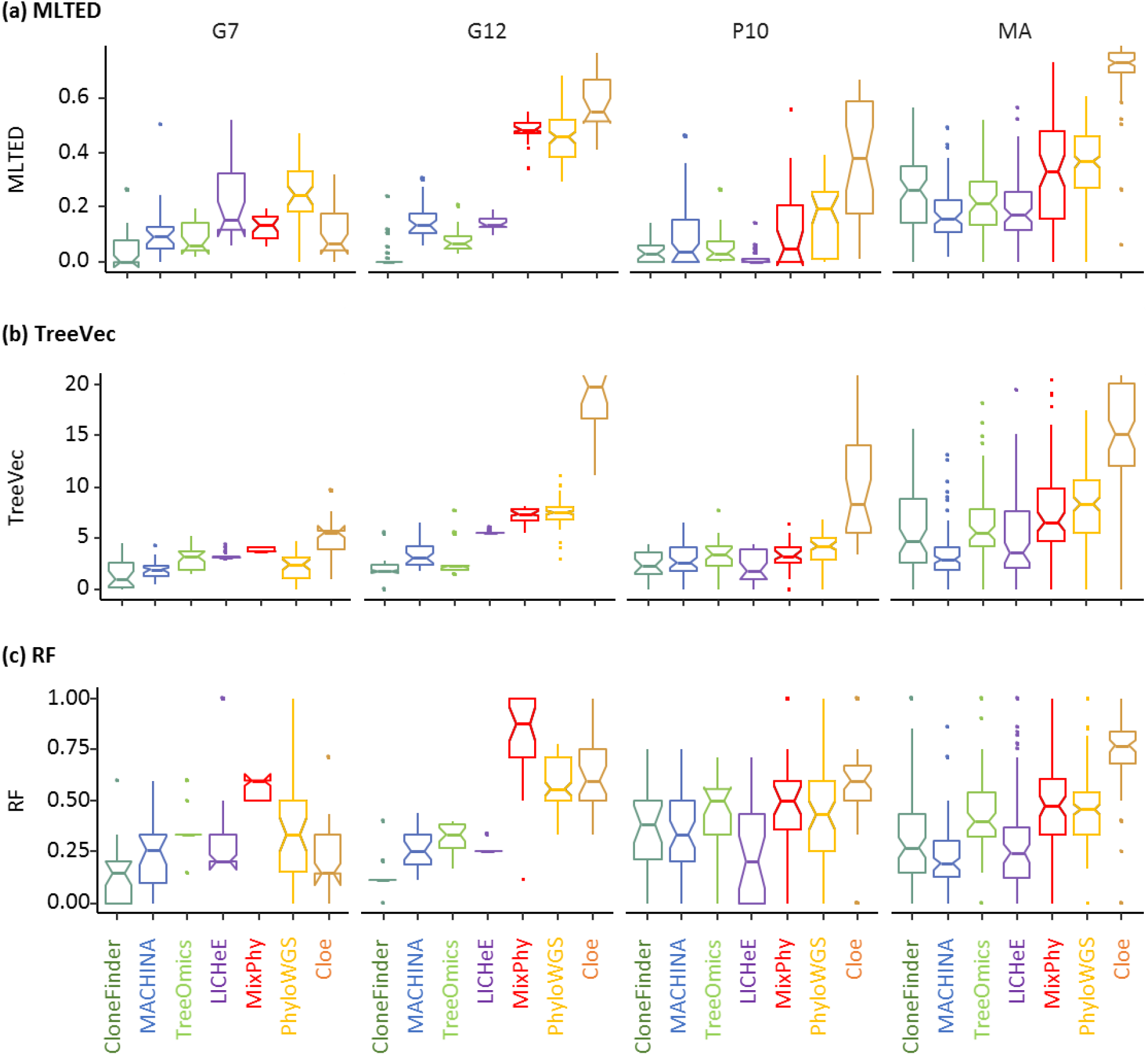
Performance of seven methods measured by (**a**) MLTED, (**b**) TreeVec, and (**c**) RF distances. MLTEDs show the accuracy of inferred mutation orders, whereas TreeVec and RF the accuracy of inferred clone phylogenies (small values indicate higher accuracy).

The clonal structures of tumors in P10 and MA datasets were more complex than G7 and G12 datasets. The P10 datasets were composed of a few tumor samples, in which ancestral clones were present alongside their descendants (**Fig. 1e** and **1f**). The MA datasets were generated by simulating the evolution of primary and metastatic tumors. The clonal structure of metastatic tumors of some MA datasets was evolutionarily complex, as more than one founding (seeding) clone migrated from another tumor site(s) (e.g., **Fig. 1g** and **1h**). For the P10 and MA datasets, we found that the MLTED scores of Cloe were higher (worse) than other methods (**Fig. 4a**). For the MA datasets, MLTED scores of all the methods were generally higher than the other datasets, and there were large differences among the datasets. Overall, MACHINA and LICHeE showed slightly better performance than the other methods.

Next, we evaluated error rates of ordering sequential, concurrent, and parallel mutations (**Fig. 3**). We generated all possible pairs of SNVs (mutations) and classified them into these three possible categories. In each category, we computed the proportion of real mutation pairs that were not present in the inferred tree, and the proportion of all incorrect mutation pairs. The average of these two proportions was used to assess the error rate of ordering the given type of mutations (see the **Methods** section for details). Sequential and concurrent mutations were inferred with lower accuracy than the parallel mutations (**Table 1**), a difference that was greater for P10 and MA datasets. For example, the error rate of inferring parallel mutations was only 4 – 6% in CloneFinder, MACHINA, Treeomics, and LICHeE analyses for MA datasets, while the error rates for sequential and concurrent mutations were much higher (12 – 21%). Therefore, identification of parallel mutations was generally more reliable than classifying sequential or concurrent mutations.

### Accuracy of predicting branching patterns (topology of clone phylogeny)

We next evaluated the accuracy of inferred branching patterns by computing TreeVec and RF distances (see the **Methods** section for details). These distances evaluate the errors of clone groupings in inferred phylogenies. For the G12 datasets (**Fig. 4b**), CloneFinder, MACHINA, Treeomics, and LICHeE showed smaller TreeVec distances than the other methods, i.e., these methods produce more accurate branching patterns. Cloe generally showed higher TreeVec distances than other methods. For the G7 datasets, all the methods showed relatively small TreeVec, and indeed, reconstructed clone phylogenies were quite similar to the correct phylogeny for these data (**Additional file 1: Fig. S1**). These patterns are consistent with those based on MLTED scores (**Fig. 4a** and **Table 1**). The results of RF distances were also consistent with MLTED and TreeVec analyses (**Fig. 4c**).

### Impact of persisting ancestral clones

To better understand factors that cause inference errors, we analyzed the impact of the presence of ancestral clones in tumor samples on the accuracy of clone inference. We found that fewer than 50% of the ancestral clones were identified by current methods (**Fig. 5**). Treeomics analysis rarely identified ancestral clones, even in datasets containing as many as six ancestral clones, and MixPhy also performed poorly.

**Figure 5:**
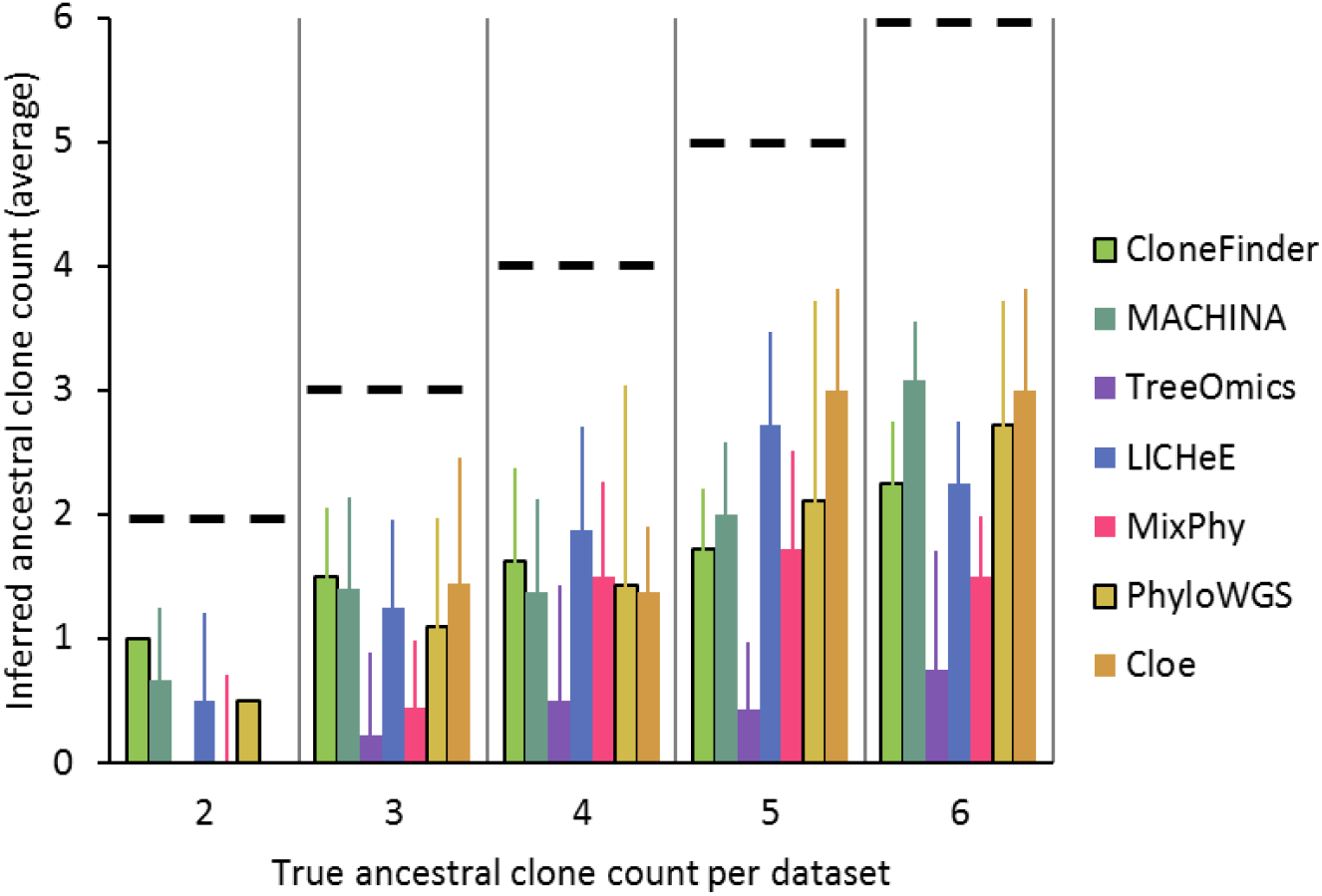
The average number of ancestral clones that were identified per dataset for the P10 datasets. We grouped P10 datasets based on the true number of ancestral clones in a dataset. For each dataset, we counted the number of ancestral clones identified by a clone prediction method. We then computed the average across the dataset. Dashed lines are the correct count. Error bars represent standard deviation values.

All tested methods, except for Cloe, performed well in ordering mutations for a dataset that contained only two ancestral clones (**Fig. 6a**). However, the accuracy of ordering mutations declined when datasets contained tumors with a large number of ancestral clones. In these datasets, CloneFinder, MACHINA, Treeomics, and LICHeE analyses generally had a lower error, indicating their robustness to the presence of persisting ancestral clones within a dataset.

**Figure 6:**
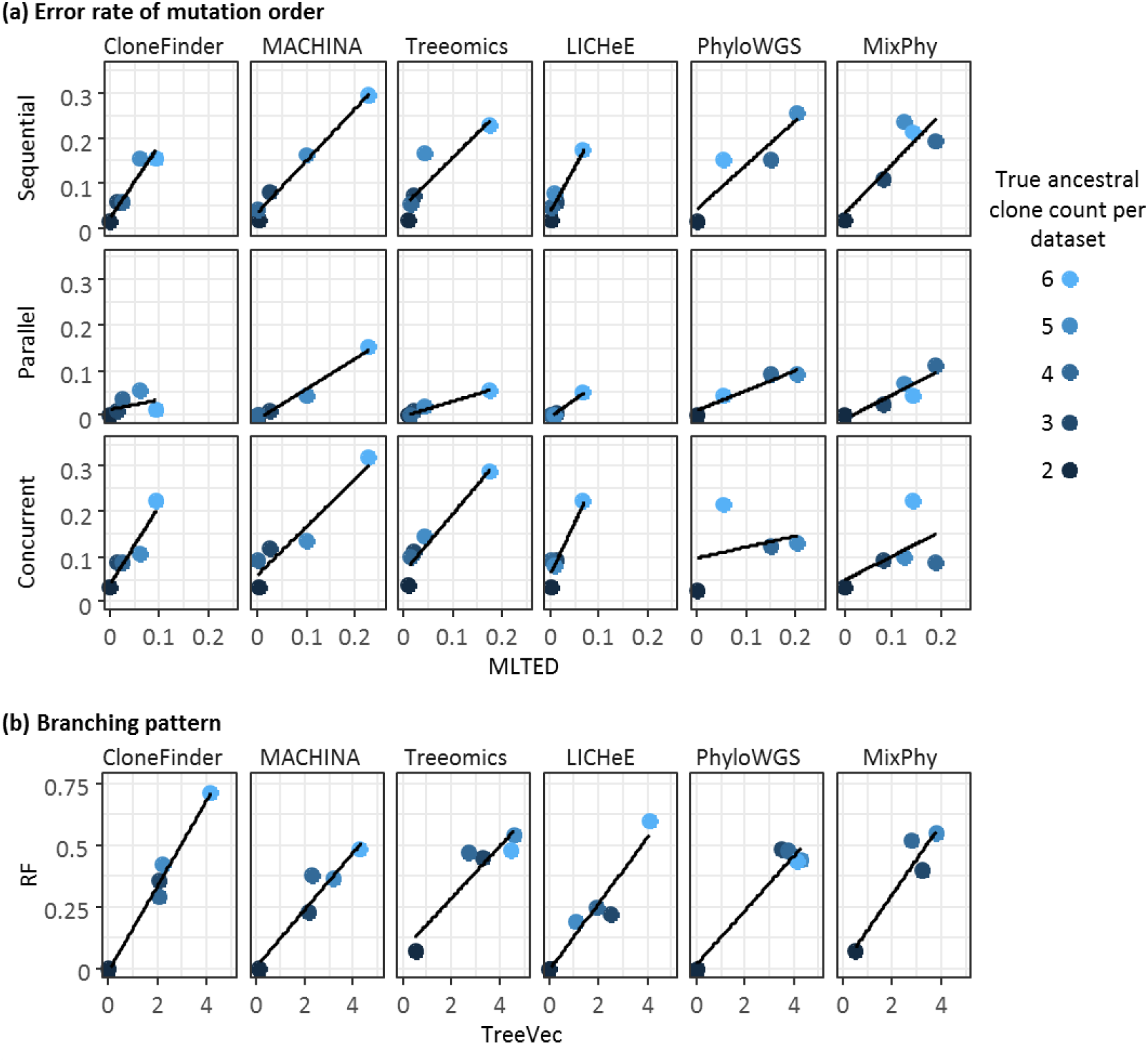
Accuracy of ordering mutations and inferring branching patterns, when a dataset contained various number of of ancestral clones. P10 datasets were used and were grouped based on the true ancestral clone count in a dataset. Each point shows the average of tree distance across the datasets. (**a**) The average error rate of ordering mutations and MLTED. (**b**) RF distances and TreeVec. Cloe method was excluded because both MLTED scores and average error rates were very high (**Fig. 4** and **Table 1**).

For Treeomics, LICHeE, and CloneFinder, the error rate of predicting parallel mutation did not increase significantly with an increasing number of ancestral clones, but the error rates in predicting sequential and concurrent mutations increased significantly (**Fig. 6a**). This is because the inability to detect ancestral clones would misclassify sequential mutations as concurrent mutations (e.g., **Additional file 1: Fig. S2**).

Consistent with the ability to predict correct mutation orders, all tested methods (except for Cloe) showed relatively small TreeVec and RF distances when a dataset contained only two ancestral clones (**Fig. 6b**), while CloneFinder, MACHINA, Treeomics, and LICHeE generally produced smaller TreeVec and RF distances for datasets with larger numbers of ancestral clones. Overall, no method produced highly accurate clone phylogenies for datasets containing a large number of ancestral clones.

### Impact of polyclonal seeding events during metastatic tumor evolution

The analysis of MA datasets was used to assess the impact of polyclonal seeding of metastatic tumors on clone phylogeny and mutation orders. These datasets contained primary tumors and four or six metastatic tumors. Up to four metastatic tumors per dataset evolved with polyclonal seeding events, i.e., these metastatic tumors were founded by more than one seeding clone. When a metastatic tumor received more than one seeding clone (polyclonal seeding events), these tumors contained clones from different evolutionary lineages due to distinct founder (seeding) clones (e.g., **Fig. 1g** and **1h**). No tested method was able to accurately identify a majority of clones within multiple-seeded metastatic tumors (**Fig. 7a**). MACHINA is the only method that incorporates the metastatic progression model of clone seeding events during its estimation process, and it did outperform other tested methods when datasets contained the largest number of multiple-seeding events (**Fig. 7a**). Overall, the poor performance of all the methods in inferring clones resulted in higher error rates of ordering mutations and reconstructing branching patterns (**Fig. 7b–7g**).

**Figure 7:**
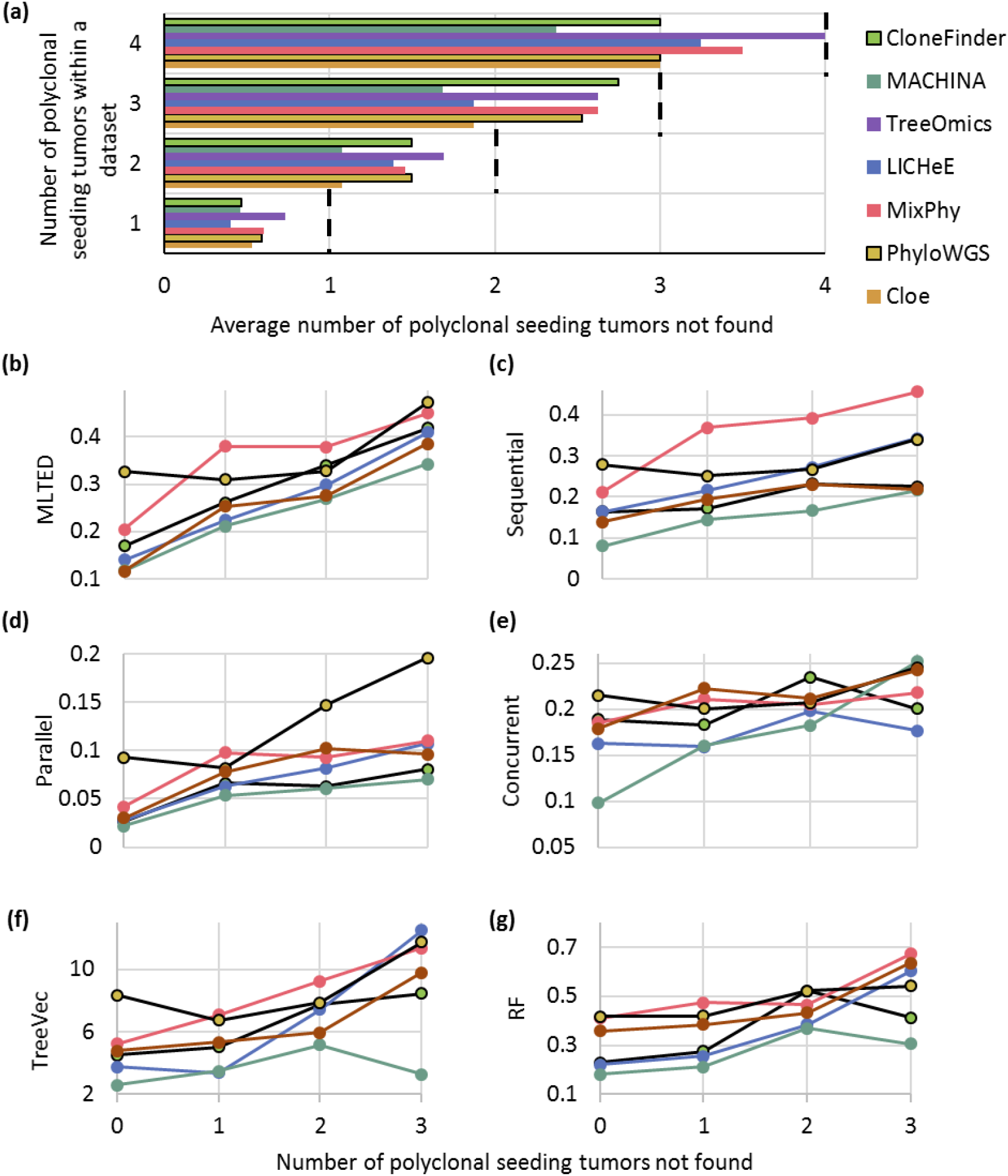
Accuracy of identifying different lineage clones within a tumor for the MA datasets. (**a**) The average count of metastatic tumors with polyclonal seeding events that were not predicted. (**b-g**) MLTED, error rates of ordering mutations, TreeVec, and RF distances. The x-axis for panels **b-e** is the same as those in panels **f** and **g**. We excluded Cloe because its MLTED score was very high, i.e., clone phylogenies were inaccurate.

Even when a MA dataset contained only one polyclonal seeding event in a metastatic tumor, we observed errors in phylogenetic predictions, mainly caused by unsuccessful inference of clones’ presence within that metastatic tumor. For example, **Figure 8** shows inferred clone phylogenies for an example dataset (**Fig. 1g** and **1h**) in which a metastatic tumor (M5) experienced polyclonal seeding events such that two seeding clones came from two distinct clone lineages (clone lineage C/D, which contained clone C and D, and lineage M with clone M). All the methods, including MACHINA, identified only one out of these two clone lineages (lineages C/D or M), with MACHINA producing two solutions (**Fig. 8b** and **8c**). The first solution contained only clone C, whereas the second solution contained only clone M. In these MACHINA phylogenies, these two clones were connected with erroneously long branches (**Fig. 1g**). Thus, those correct clones found within the M5 metastatic tumor were convoluted into one clone genotype in the inferred clone phylogenies. This same type of error was observed in predicted clone phylogenies generated via other methods (**Fig. 8**), except for Cloe (which produced phylogenies that dramatically differed from the true phylogeny). Apart from these errors, the predicted clone phylogenies were largely similar to the true clone phylogeny, and the branching patterns were mostly correct (**Fig. 1** and **Fig. 8**). For this example MA dataset, MACHINA, CloneFinder, and LICHeE produced more accurate clone phylogenies than other methods. For example, Treeomics, PhyloWGS, and MixPhy produced much smaller phylogenies, as these methods did not infer many ancestral or highly-similar clones.

**Figure 8.**
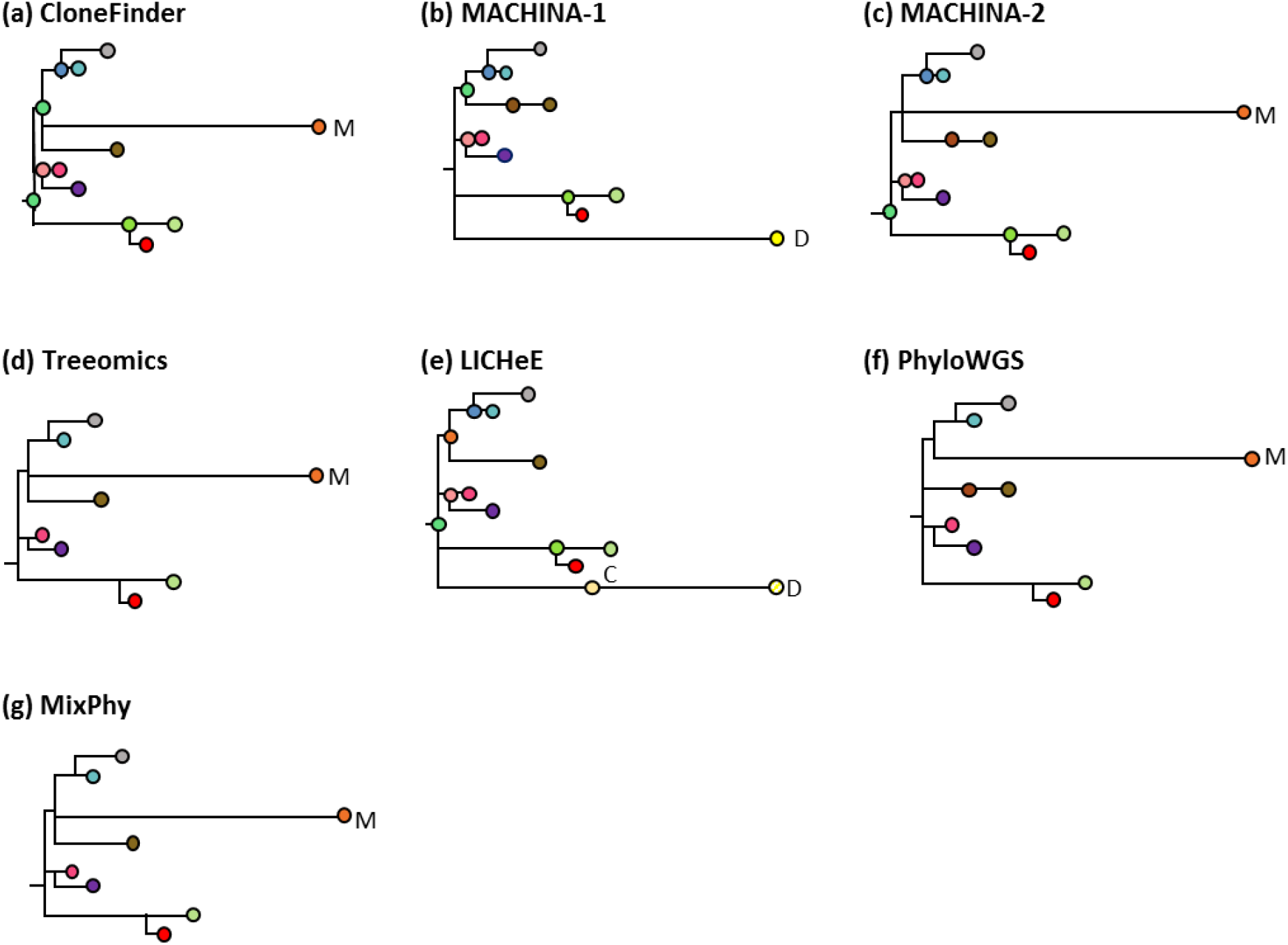
Clone phylogenies inferred by six methods (Cloe method was excluded due to error-prone) on an MA dataset. True clone phylogeny is given in **Figure 1g**. MACHINA produced two solutions (**b** and **c**). Inferred clones are annotated, and colors correspond to clones in **Figure 1g**. All the method produced either clone lineage M or lineage C/D, which were found in the M5 tumor (**Fig. 1h**). The first solution of MACHINA (**b**) produced clone D, and LICHeE produced clones C and D (**e**). The other methods produced clone M.

This pattern of errors in inferred clone phylogenies became more acute when a dataset included many metastatic tumors that evolved with polyclonal seeding events. For example, when a dataset was composed of four metastatic tumors with polyclonal seeding events, inferred clone phylogenies contained fewer clones than the true phylogeny (**Additional file 1: Fig. S3**). The tested methods tended to predict only one clonal lineage for each of the four metastatic tumors of this dataset. Note that Cloe produced a phylogeny with little similarity to the true phylogeny. MACHINA produced 870 solutions for this example dataset, with the best solution (smallest number of SNV assignment errors per clone) similar to the true phylogeny, and the worst solution that missed many clonal lineages. Overall, current clone prediction methods cannot reliably decompose many clones within metastatic tumors with polyclonal seeding events.

### Empirical data analysis

The application of these clone prediction methods to an empirical dataset (A7 dataset from a previous study [30]) showed results consistent with our analyses of simulated data. The original study reported that metastatic rib and lung tumors harbored clones from different clonal lineages (**Fig. 9a**). The lung tumor contained three different clone lineages, indicating a complicated history of metastatic tumor evolution. Different methods predicted clone phylogenies that showed limited similarity to the clone phylogeny reported in the original study (**Fig. 9b–9i**). MACHINA produced four similar solutions (**Fig. 9b–9e**). However, only the predicted evolutionary relationship of clones from liver and kidney tumors agreed with those reported in the original study [30]. The predicted clone sharing between lung and brain tumors reported by CloneFinder agreed with the original study, but the clone phylogeny differed dramatically (**Fig. 9f**). Treeomics correctly predicted the evolutionary relationship of clones from the liver, kidney, and rib tumors, but did not predict most of the ancestral clones (**Fig. 9g**), a failing that we also observed in our simulation results. PhyloWGS produced two distinct but highly similar clone phylogenies (**Fig. 9h** and **9i**) that indicated the presence of three clonal lineages, instead of the two lineages reported in the original study. LICHeE analyses did not produce a solution. MixPhy produced >400 clones for this dataset, and Cloe results suggested the unlikely scenario that all predicted clones were present in most of the samples. Therefore, we anticipate that the application of different computational methods in actual empirical data analysis will result in widely varying inferences, making it challenging to reach reliable biological conclusions, when the tumor evolution is highly complex.

**Figure 9.**
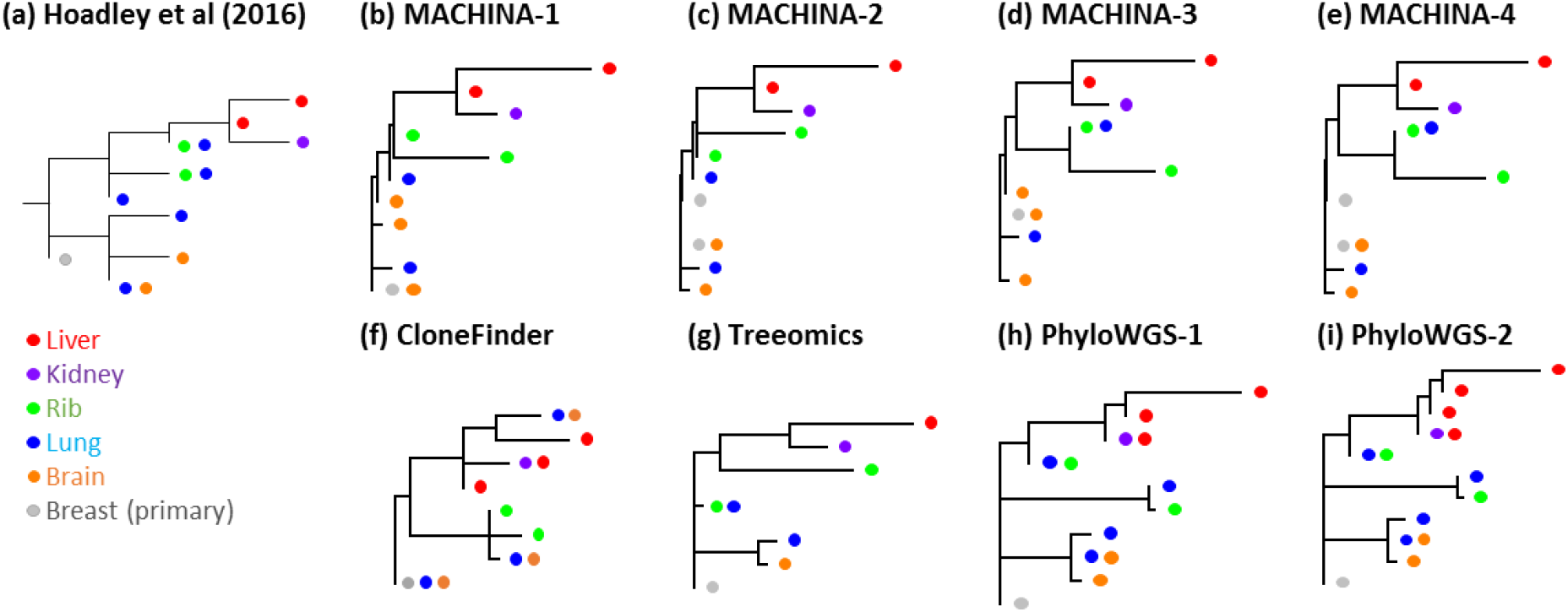
Empirical data analysis for the A7 dataset. The color of clones in the phylogeny corresponds to the location of clones’ samples. (**a**) Clone phylogeny reported by Hoadley et al. (2016). (**b-i**) Inferred clone phylogenies by using (**b-e**) MACHINA, (**f**) CloneFInder, (**g**) Treeomics, and (**h** and **i**) PhyloWGS. MACHINA and PhyloWGS produced more than one phylogeny.

## Discussion

Predictions of accurate clone phylogenies are essential to infer the order of driver mutation occurrences and the evolutionary relationship of clones. We tested the accuracy of published methods in reconstructing clone phylogenies as a first step in identifying the patterns of errors in clone phylogeny inference, which revealed some useful guidelines for applying computational methods in practical data analysis. To begin with, we suggest the use of CloneFinder, MACHINA, Treeomics, and LICHeE, because they often showed lower error rates of ordering mutations and inferring phylogenies. All of these methods benefit from the use of intrinsic evolutionary relationship of tumor clones. The evolutionary information provides resolution beyond inferences primarily based on the dissimilarities of observed SNV frequencies because low read depth cause SNV frequencies to have significant variance and clone predictions based on only the similarities of observed SNV frequencies become error-prone.

Careful consideration of the input data is strongly recommended before choosing a method for analyses. First, these clone prediction methods require copy-number-neutral SNVs, because observed SNV frequencies are affected by copy number alterations (CNAs). SNV frequencies should be adjusted to eliminate the impact of CNAs. Notably, Cloe [39] is designed for the analysis of datasets with CNAs, but it did not perform well for datasets without CNAs.

Also, most methods are known not to be robust to the presence of incorrect SNV assignments, so one should proceed with extreme caution when analyzing datasets with high rates of sequence error. For example, LICHeE may fail to produce any inferences on such datasets or the accuracy may become much lower than other methods (e.g., Treeomics) [35]. LICHeE failed to produce any results for our example empirical dataset [30]. In general, clone predictions are expected to become more challenging when the dataset contains CNAs and sequencing errors. Also, the accuracy of clone phylogeny inference can be adversely impacted by biological factors (e.g., the impact of strong natural selection).

Another important consideration in experimental design is the benefit of sequencing a larger number of tumor samples. The most successful methods in our evaluations use the intrinsic evolutionary relationships among tumor samples so a larger sample number can provide more information to improve clone predictions. All of the methods tested here performed well on simulated datasets with the largest number of tumor samples (G12 datasets). Although the actual number of tumor samples preferred depends on the situation, it is clear that one should avoid datasets generated from only a few samples. Importantly, datasets with a very small number of samples will underestimate the genetic heterogeneity of a tumor site, and therefore, the use of a large number of samples per patient is a standard recommendation [6, 45].

We do not expect any of the currently available methods to be effective in situations where each tumor sample contains clones from many lineages (if tumors frequently exchange clones). We have previously documented that CloneFinder will not perform well on such datasets [35]. Also, our simulation analyses have shown that none of the tested methods perform well when a mixture of clones from different evolutionary lineages exist within metastatic tumors (multiple seeding).

Lastly, we suggest using multiple methods to infer clone phylogenies and examining the consistency among the results. We found that the best performing methods produced similar results when inferred clone phylogenies were accurate. When using Treeomics, it is crucial to be aware that the inferred clone phylogenies will not include most of the ancestral clones. Also, potential errors on clonal lineage deconvolution can be detected when MACHINA produces at least two disparate clone phylogenies (e.g., **Fig. 8**) or when MACHINA produces hundreds of solutions. In general, the inconsistency of inferred clone phylogenies suggests the influence of complicated clonal structures within tumors, i.e., a mixture of different lineage clones. Currently, no method can produce accurate clone phylogenies from such data. Thus, consistency among inferred phylogenies may be useful to validate inferences.

In summary, we can accurately infer clone phylogenies only when tumor evolution generally tracks clonal evolution, a relationship that is disrupted when tumors exchange clones reduce the quality of the inferred clone phylogenies. Also, ancestral clones that persist alongside the descendant clones within a tumor sample are difficult to identify, leading to inaccuracy in the reconstruction of evolutionary events.

## Conclusions

Analyses of correct clone phylogenies are critical to a better understanding of tumor evolution and the influence of genetic heterogeneity. We recommend clone prediction methods that use the intrinsic evolutionary relationship of tumor samples (e.g., CloneFinder, MACHINA, TreeOmics, and LICHeE). The inferences of multiple methods should also be compared to validate predictions. There is a strong need for more advanced methods that can perform well for datasets with intermixing of tumor samples.

## Methods

### Generation of bulk sequencing data

We analyzed 150 simulated datasets that were available from published studies in which the accuracy of inferred clone sequences was assessed [13, 35]. Each dataset contained information on mutant and wild-type read counts (with read counting errors).

#### G7 and G12 datasets

These datasets contained seven and twelve clones, respectively, modeled after the predicted evolutionary histories of two patients (EV005 and RK26 [44], respectively) (**Fig. 1a–1d**) [35]. Each tumor sample may contain one or a few evolutionarily closely-related clones, assuming a localized genetic heterogeneity due to branching evolution [4, 6]. Thus, the migration of cancer cells to another section of a tumor was assumed to be rare in these datasets. In total, we obtained 60 simulated datasets (replicates) with 34-89 SNVs per dataset.

#### P10 datasets

In these datasets, various numbers of clones persisted within a sector (sample) of a tumor after the origin of descendant clones. Ten random clone phylogenies were simulated, and every tumor sample was populated with one tip clone and its ancestral clones (“localized sampling process” [34]) (**Fig. 1e** and **1f**). Each of P10 datasets contained 2 – 6 ancestral clones (30 datasets). A selection of simulated clone phylogenies is shown in Figure 3 of Miura et al. [35].

#### MA datasets

These datasets were generated by modeling the evolution of primary and metastatic tumors (four or seven metastatic tumors per dataset) [13]. Metastatic tumors were founded by cancer cells (seeding clones) that migrated from another tumor site (primary or another metastatic tumor). Under a simple metastatic tumor evolution scenario, each metastatic tumor received a single founder (seeding) clone from another tumor site, and a metastatic tumor contained only clones that evolved from a single seeding clone. Clonal structures of metastatic tumors became more complicated when a metastatic tumor was seeded by more than one clone (polyclonal seeding events). In MA datasets, a metastatic tumor received a maximum of two seeding clones, and any dataset may contain more than one metastatic tumor with polyclonal seeding events. Thus, the observed genotypes of these metastatic tumors represented two convoluted clone lineages, and clone prediction methods were required to correctly identify such tumors and decompose them into two distinct clone lineages (e.g., **Fig. 1g** and **1h**). Each MA dataset contained up to four metastatic tumors with polyclonal seeding events. Each clone phylogeny was unique (60 MA datasets). All the clone phylogenies are shown in **figure 2**.

### Selection of clone prediction methods and parameter settings

We selected clone prediction methods that have performed well in predicting clone genotypes from observed SNV frequencies or read counts of bulk sequencing data [35]. That is, we excluded methods that produce highly incorrect clone genotypes because such clone genotypes do not produce correct clone phylogenies. By this criterion, we excluded CITUP [46], BayClone2 [47], Clomial [48], Canopy [49], cloneHD [50], and AncesTree [51] (see **Additional file 1: Table S1** for the average number of SNV assignment errors per clone). We did not include methods that require prior information of the composition of SNV clusters (e.g., TrAp [52]) or those that require the use of another software to produce clone genotypes by ordering predicted clusters (e.g., PyClone [53] and SciClone [54]). Lastly, we did not include methods that were designed for the analyses of single-cell sequencing data (e.g., SCITE [55] and BEAM [56]), because clone deconvolution is not necessary for this type of data, while these methods focus on imputing missing data and minimizing SNV assignment errors in the inference of cell phylogenies [31, 32]. These considerations resulted in the selection of seven clone prediction methods (**Table 1**) [13, 34–39]. Each method was used with its default or recommended parameter settings. In MA datasets, we found many similar clone genotypes, so we used parameter settings that can differentiate similar clone genotypes. This modification was applied only for LICHeE and CloneFinder, as only these two methods include options for this purpose.

#### MACHINA [13]

We used the PMH-TI mode in the MACHINA software, which infers clone genotypes from read count data. The MACHINA software requires *a priori* identification of tumor sites as primary or metastatic for each sample. Since G7, G12, and P10 datasets were simulated without the consideration of primary and metastatic tumor evolution, we assumed that the primary tumor contained the root clone (e.g., clone A for G7 and G12 datasets) (**Fig. 1a and 1c**). When a root clone was not present in a dataset, we selected the clone that was most closely located to the root of a simulated phylogeny. For MA datasets, we provided the correct tumor site (primary or metastatic site, in which distinct metastatic tumor sites were accordingly distinguished) for each clone sequence that was found. Note that MACHINA often produced a large number of solutions (>10 solutions per dataset) for G7, G12 and MA datasets. In those cases, we first identified the best and worst solutions for each dataset, which were determined based on the average number of SNV assignment errors per clone. We reported the average error rate (see below) of the best and worst solutions.

#### LICHeE [34]

Following the default settings, we set the variant allele frequency (VAF) error margin the value 0.1. SNVs were considered robustly present in a sample at VAF > 0.005 (robust SNVs), and the others were considered absent in a sample. SNVs with VAF > 0.6 were excluded. LICHeE groups SNVs based on the pattern of presence/absence of mutations across the samples and each SNV group was required to contain at least two robust SNVs. LICHeE also clusters SNVs by VAF similarities. We required that an SNV cluster contained at least two SNVs unless an SNV was sample specific. All the SNV groups/clusters were initially kept in the network. Two groups/clusters could collapse when mean VAF difference was < 0.2.

LICHeE did not produce clonal compositions of samples (i.e., clone frequencies). Thus, we estimated clone frequencies using the relationship ½*f* × M = *V*, where *f* is a two-dimensional matrix of estimated clone frequencies of the samples, M is a matrix of predicted clone genotypes, and *V* is the observed SNV frequency [33]. The equation above applies to cases where the variants are free of copy number alterations (CNAs) [33], which is the case for our datasets. We estimated *f* through the regression of *V* to a function of M and *f* [57]. Clone frequencies were estimated excluding SNVs with small total read count (<50) and mutant read count (<2), because those observed SNV frequencies were not reliable. When ancestral clones were predicted to co-exist with their descendant clones within a sample, we tested if these ancestral clones were spurious. Between a pair of ancestral and descendant clones, we compared observed SNV frequencies that are unique to the descendant clone and those shared with the ancestral clone. We used the expectation of higher observed SNV frequencies on shared (mutations that were found in both clones) than on unique mutations (mutations that were found in only a descendant clone; *t*-test) to discover the spurious presence of ancestral clones. When the differences between SNV frequencies were not significant (*P* > 0.05), the ancestral clones were removed. Also, we discarded clones present at low frequencies (<2%).

In the analyses of MA datasets, only SNVs with zero SNV frequency were considered to be robustly absent from a sample, and SNVs with > 0.0001 frequency were considered to be robustly present in a sample (robust SNVs). All SNVs were examined regardless of their observed frequency. The minimum number of SNVs per cluster/group was set to one. Two SNV clusters were collapsed when mean SNV frequency differences were less than 1%. We did not discard any ancestral clones.

#### CloneFinder [35]

We estimated clone genotypes using SNVs with at least 50 reference read counts and two mutant read counts, and we discarded clones when estimated clone frequencies were < 2%. To analyze MA datasets, we did not combine similar clone genotypes or discard clones. We used all reads.

#### Treeomics [36]

We used the option of enabling subclone detection.

#### PhyloWGS [37]

The fraction of expected reference allele sampling from the reference population and the variant population were 0.999 and 0.4999, respectively. We set copy number equal to one (heterozygous mutant allele). As PhyloWGS did not produce clone frequencies, we computed clone frequencies using the approach described for LICHeE (see above).

#### Mixed Perfect Phylogeny (MixPhy) [38]

We performed analyses in MixPhy (v0.1) with the option of a heuristic algorithm. As the input file requires a binary matrix of tumor sample genotypes (presence/absence of mutation), we provided correct sample genotypes, assuming that there were no false positive or false negative detections of mutations.

#### Cloe [39]

We applied Cloe with 10,000 iterations and 4 MCMC parallel chains at temperatures of 1, 0.9, 0.82, and 0.75. For the posterior evaluation of MCMC sampled trees, the burn-in of MCMC chains was 0.5, and chain thinning was 4. The maximum number of clones for a dataset was set to the true clone count.

### Evaluation of predicted clone phylogenies

We compared each predicted clone phylogeny with the respective true clone phylogeny by using the following four metrics.

#### Multi-labeled tree edit distance (MLTED) [58]

A clone phylogeny is often viewed as a mutational tree [43] in which all the mutations are mapped along branches. Mutational trees are useful when the number of tips in the inferred clone phylogeny differs from the true phylogeny and when the sequences of the inferred clones do not match all the true clones. We used the Multi-labeled Tree Edit Distance (MLTED score) for comparing the inferred and the true tree, as it has been designed to evaluate clone trees [58], available at https://github.com/khaled-rahman/MLTED. This algorithm requires that the inferred tree contains the same set of mutations as in the true tree. Because of errors in clone sequence predictions, some mutations were not assigned to any branch in the inferred tree. These mutations were placed at the root of the inferred mutational tree.

#### The error rate of ordering mutations

We generated all possible pairs of SNVs (mutations) and classified them into three possible types, i.e., concurrent, sequential, and parallel (see **Fig. 3** for examples). Concurrent mutations are those that occurred on the same branch (irrespective of their order), whereas sequential and parallel mutations are those that occurred on different branches of the clone phylogeny. More specifically, two mutations are sequential if one occurred on the ancestral branch and the other on its descendant branch, but multiple intervening branches may separate them. Two mutations are parallel if they are found on sibling lineages that have descended from their most recent common ancestor. Any true mutation pair not found in the inferred tree was classified as “unassigned.”

We estimate the error rate of ordering concurrent, sequential, and parallel mutations, separately. In each category, we first scored the number of true mutation pairs that were not present in the inferred tree and divided it by the total number of true mutation pairs. Then, we scored the number of mutation pairs that were incorrect and divided it by the total number of inferred mutation pairs. Then, the average of these two proportions was used as the error rate of ordering the given type of mutations. Similar measures have been used to evaluate clone prediction methods in previous studies [34].

#### Advanced Tree vector (TreeVec) [59]

We also evaluated the accuracy of branching patterns (topology) in inferred clone phylogenies (clonal lineage trees [43]). For this purpose, we first mapped inferred clone genotypes to the true clone genotypes, because inferred clone genotypes never perfectly match the true clone genotypes. We mapped each inferred clone genotype to its most similar true clone genotype in a two-step process. First, each true clone genotype was compared to all the inferred clone genotypes, and the two clones with the smallest difference were paired. When the number of inferred clones was greater than the number of true clones, the remaining inferred clones were paired with the most similar true clone genotype. For uniformity, we reconstructed inferred clone phylogenies by using predicted clone genotypes produced by each method. Because mutations arose only once in the computer simulated data, the maximum parsimony analysis was suitable [60] and was performed using MEGA-CC [61]. All the clone phylogenies were rooted using germline sequences (normal cells) as outgroups. In inferred clone phylogenies, we labeled tips with clone annotations. When an inferred clone genotype had two different annotations, we duplicated the genotype in an inferred clone phylogeny, i.e., the corresponding tip was duplicated. Also, two inferred clone genotypes might have the same annotation. In this case, two tips in an inferred clone phylogeny were labeled identically.

Among various tree distance computation methods for phylogenies [62], we selected the advanced TreeVec distance developed by Kendall et al. [59], because TreeVec allowed more than one tip with identical labels. Briefly, TreeVec distance computation first collapsed any monophyletic clade(s), i.e., a clade with tips that had an identical label. Then, the traditional TreeVec distance [63] was computed, which counted the number of branches (edges) between the root and the node of the most recent common ancestor (MRCA) of a pair of clones. For all pairs of clones, the Euclidean metric between inferred and true counts was computed. We used the treespace software [64] to compute this advanced TreeVec distance.

#### Robinson and Foulds (RF) distance [65]

We also computed RF tree distance, because it is widely applied in the evaluation of species phylogenies. We used PhyloNET software [66] to count the number of partitions that were common and different between the true and the inferred phylogeny. The RF distance is the number of differing partitions divided by the total number of partitions in the two phylogenies. Note that RF distance computation requires that both the inferred and the true clone phylogenies contain the same number of tips (clones). However, inferred clone phylogenies may contain more tips than the respective true phylogenies, when more than one tip is assigned an identical clone annotation (i.e., more than one inferred clone genotype was similar to a true genotype). When there were too many tips in the inferred tree, we retained only those tips that showed the highest similarity to the true clone genotypes, such that each true clone genotype was matched with exactly one inferred genotype.

### Empirical data analyses

We obtained an empirical dataset (patient A7 dataset [30]; https://github.com/raphael-group/machina), which contained genotypes for 478 copy-neutral SNVs. This dataset contains SNV frequencies of one primary tumor sample (breast) and four metastatic tumors (lung, liver, rib, and brain), for which clone phylogenies and clonal composition of each sample have been previously reported [30]. For real data, true clone genotypes are not available, so we annotated each clone on the inferred phylogeny based on the sample(s) that contained it (Fig. 9a) in order to compare the reported phylogeny [30] with those inferred by the clone prediction methods listed in **Table 1**.

## Abbreviations

SNV: single-nucleotide variant
VAF: variant allele frequency
CNV: copy number alteration

## Declarations

### Ethics approval and consent to participate

Not applicable

### Consent for publication

Not applicable.

### Availability of data and material

The G7, G12, and P10 datasets are available on the website of the CloneFInder software (https://github.com/gstecher/CloneFinderAPI). The MA datasets and A7 dataset were obtained from the website of the MACHINA software (https://github.com/raphael-group/machina).

### Authors’ contributions

SK conceived the project. SM and SK designed analyses. SM, TV, JD, TB, and JC performed the analyses. SK, SM, and TV wrote the manuscript. All authors read and approved the final manuscript.

## Acknowledgments

We thank Drs. Antonia Chroni, Heather Rowe, Louise A Huuki, Zachary Hanson-Hart, and Viriya Keo for critical comments and technical support.

## Additional files

Additional file 1: **Table S1 and Figures S1-S3**. Supplementary table and figures. (PDF)

